# SHP2 Is Required for BCR-ABL1-Induced Hematologic Neoplasms

**DOI:** 10.1101/157966

**Authors:** Shengqing Gu, Azin Sayad, Gordon Chan, Wentian Yang, Zhibin Lu, Carl Virtanen, Richard A. Van Etten, Benjamin G. Neel

**Affiliations:** Department of Medical Biophysics, University of Toronto, Toronto, ON M5G1L7, Canada; Princess Margaret Cancer Center, Toronto, ON M5G2M9, Canada; Department of Orthopaedics, Brown University Alpert Medical School, Providence, RI 02912, USA; Chao Family Comprehensive Cancer Center, Division of Hematology/Oncology, University of California, Irvine, Irvine, CA 92697, USA; Present address: Department of Biostatistics and Computational Biology, Dana-Farber Cancer Institute, Boston, MA 02215, USA; Current address: Laura and Issac Perlmutter Cancer Center, NYU Langone Medical Center, New York, NY 10016, USA

## Abstract

BCR-ABL1-targeting tyrosine kinase inhibitors (TKIs) have revolutionized treatment of Philadelphia chromosome-positive (Ph^+^) hematologic neoplasms. Nevertheless, acquired TKI resistance remains a major problem in chronic myeloid leukemia (CML), and TKIs are less effective against Ph^+^ B-cell acute lymphoblastic leukemia (B-ALL). GAB2, a scaffolding adaptor that binds and activates SHP2, is essential for leukemogenesis by BCR-ABL1, and a GAB2 mutant lacking SHP2 binding cannot mediate leukemogenesis. Using a genetic loss-of-function approach and bone marrow transplantation (BMT) models for CML and BCR-ABL1^+^ B-ALL, we show that SHP2 is required for BCR-ABL1-evoked myeloid and lymphoid neoplasia. *Ptpn11* deletion impairs initiation and maintenance of CML-like myeloproliferative neoplasm, and compromises induction of BCR-ABL1^+^ B-ALL. SHP2, and specifically, its SH2 domains, PTP activity and C-terminal tyrosines, is essential for BCR-ABL1^+^, but not WT, pre-B cell proliferation. The MEK/ERK pathway is regulated by SHP2 in WT and BCR-ABL1^+^ pre-B cells, but is only required for the proliferation of BCR-ABL1^+^ cells. SHP2 is required for SRC family kinase (SFK) activation only in BCR-ABL1^+^ pre-B cells. RNAseq reveals distinct SHP2-dependent transcriptional programs in BCR-ABL1^+^ and WT pre-B cells. Our results suggest that SHP2, via SFKs and ERK, represses *MXD3/4* to facilitate a MYC-dependent proliferation program in BCR-ABL1-transformed pre-B cells.

## Introduction

The Philadelphia (Ph^+^) chromosome translocation t(9, 22) generates the oncogene *BCR-ABL1*, encoding a constitutively activated tyrosine kinase. Different BCR-ABL1 isoforms cause chronic myeloid leukemia (CML) and Ph^+^ B-cell lymphoblastic leukemia (B-ALL).1,2 ABL1 tyrosine kinase inhibitors (TKIs), such as imatinib, have revolutionized the treatment of Ph^+^ hematopoietic neoplasia. Nevertheless, >40% of CML patients are PCR-positive after 10 years of imatinib treatment and remain at risk of relapse,^3^ perhaps because CML stem cells (CML-SCs) are insensitive to BCR-ABL1 inhibitors.^4–6^ The response to TKIs (combined with chemotherapy) is poorer in Ph^+^ B-ALL,^7^ with several TKI resistance mechanisms proposed, including *BCR-ABL1* mutation/amplification, elevated drug exporters, and upregulation of other oncogenic pathways.^8–10^ Therefore, new approaches are needed to eradicate BCR-ABL1^+^ neoplasia.

CML-like MPN can be reproduced in mice by retroviral transduction of *BCR-ABL1* into hematopoietic stem cell (HSC)-enriched bone marrow (BM) cells in the presence of myeloid cytokines, followed by transplantation into irradiated recipients.^11–13^ B-ALL can be induced by transducing bulk BM cells in the presence of interleukin-7 (IL-7) before transplantation.^12^ In Ph^+^ cell lines and mouse leukemia models, BCR-ABL1 is phosphorylated on Y177, which recruits the adaptor GRB2 and, thereby, the scaffolding adaptor GAB2.^14,15^ Consequently, GAB2 is constitutively tyrosyl-phosphorylated and binds SHP2 and the p85 subunit of PI3K to activate the MEK/ERK and PI3K/AKT pathways, respectively.^16,17^ Y177F mutation compromises myeloid transformation and leukemogenesis,^18–20^ and GAB2 is required both for BCR-ABL1-induced myeloid and lymphoid leukemogenesis.^21^ At present, it is not feasible to pharmacologically target GAB2, rendering it essential to identify and validate GAB2-interacting proteins that mediate leukemogenesis. Reconstituting Gab2^-/-^ donor cells with a GAB2 mutant lacking its SHP2 binding sites does not restore myeloid or lymphoid leukemogenesis, suggesting that SHP2 is required for these diseases.^21^ Nevertheless, the functions of SHP2 are not mediated solely through GAB2, and its role in BCR-ABL1-induced neoplasia remains undefined.

SHP2, encoded by *PTPN11,* is a ubiquitously expressed non-receptor protein-tyrosine phosphatase (PTP) required for full RAS-ERK pathway activation in response to growth factors and cytokines. SHP2 also modulates the PI3K-AKT and JAK-STAT pathways.^17,22–25^ SHP2 catalytic activity is suppressed by intra-molecular interaction of its N-SH2 and PTP domains, preventing substrate binding and catalysis.^26,27^ Following agonist stimulation, auto-inhibition is relieved by binding of pY peptides (such as in phospho-GAB2) to the SHP2 SH2 domains, leading to altered subcellular localization, PTP domain exposure, and enzyme activation.^17,23,26–29^

SHP2 has critical roles in hematopoiesis and leukemogenesis. Myeloid and erythroid differentiation are impaired in embryonic stem cells (ESCs) expressing mutant SHP2.^30^ SHP2 deficiency leads to defective growth factor- and cytokine-evoked ERK and AKT activation, resulting in impaired self-renewal and apoptosis of adult HSCs.^31,32^ Moreover, somatic gain-of-function alleles of *PTPN11* cause >30% of juvenile myelomonocytic leukemia (JMML) cases, are found in ~5% of AML and ALL patients, and can cooperate with *Tet2* deficiency and *AML1-ETO* to generate AML in mice.^33–37^

Multiple studies implicate SHP2 in BCR-ABL1-induced pathogenesis. SHP2 is constitutively phosphorylated in BCR-ABL1-transformed cells,^38,39^ interacts with GAB2,^16,21^ and is required for BCR-ABL1-evoked transformation of a yolk sac cell line.^40^ However, the role of SHP2 in adult Ph^+^ hematopoietic neoplasia remains elusive. Here, we utilize mouse models to address this issue, report a critical role for SHP2 in myeloid and lymphoid Ph^+^ neoplasia, and uncover a differential requirement for SHP2 in normal versus leukemic pre-B cells.

## Materials and Methods

### Mice

*Ptpn11^flox/flox^* mice^41,42^ were bred to *Mx1-Cre* or *Rosa26-creERT2* mice (The Jackson Laboratory) in the C57BL/6 background. Genotyping was performed as described.^31^

### Virus production

Replication-defective ecotropic retroviral stocks of BCR-ABL1-expressing p210MIGFP, p210MIGFPCre, and p210MINVneo^16,43,44^ were generated by transient transfection of 293T cells.^21^ Viral supernatants were collected 48 and 72 hours post-transfection and stored at −80°C.

### Mouse models of CML and B-ALL

For CML-like MPN, BM was flushed from femurs and tibiae. Red blood cells (RBCs) were lysed in 0.16M NH_4_Cl, and RBC-depleted BM cells were incubated with rat anti-mouse lineage (Lin) antibodies (CD3, CD19, Gr1, and Ter119 (BioLegend) and CD4, CD8α, CD127, and B220 (eBioscience)) for 30 minutes. Lin^+^ cells were depleted with sheep anti-rat Dynabeads (Invitrogen) for 1 hour, and the remaining cells were pre-stimulated overnight in IMDM-15%FBS, supplemented with IL-3 (6ng/ml), IL-6 (10ng/ml), and SCF (20ng/ml). On each of the following two days, pre-stimulated Lin^-^ cells were spin-infected, and on the third day, cells were harvested, resuspended in cold PBS, and injected intravenously (IV) into 6-Gy irradiated syngeneic recipients.^45^

For B-lymphoid leukemogenesis, RBC-depleted BM was resuspended in IMDM-15%FBS, supplemented with IL-7 (10ng/ml), and infected as above. After infection, cells were cultured at 37°C for 4 hours, resuspended in cold PBS, and injected IV into 11-Gy irradiated syngeneic recipients.^12^

### B-lymphoid progenitor cultures

RBC-depleted BM cells were incubated for 30 minutes with rat anti-CD4, -CD8, -Gr1, -Mac-1 and -Ter119 antibodies, followed by sheep-anti-rat Dynabeads for 1 hour. Following magnetic separation, the remaining cells were cultured in 24-well plates in OptiMEM-10%FBS containing 5ng/ml IL-7 and 50µM β-mercaptoethanol.^46^

### Flow cytometry

All studies used an LSRII flow cytometer. RBC-depleted cells from peripheral blood, spleen, and/or bone marrow were labeled with antibodies to myeloid (Gr-1, Mac-1), B lymphoid (B220, CD19), and T lymphoid (CD4, CD8) markers. To assess apoptosis, cells were washed in cold PBS, resuspended in Annexin V staining buffer (BD Biosciences), and incubated with Annexin V (PE- or FITC-conjugated; 1:300) and Sytox Blue (1:1000) for 20 minutes at room temperature in the dark. Samples were analyzed within 1 hour, with low-FSC apoptotic bodies gated. For cell cycle analysis, cells were resuspended in OptiMEM-10%FBS, supplemented with 5 ng/ml IL-7 containing 10 µM Hoechst-33342. After 30 minutes at 37°C, Pyronin-Y was added (2.5 µg/ml), and cells were incubated for another 15 minutes prior to analysis.

### Immunoblotting

Myeloid- or B lymphoid-enriched BM cells were transduced with p210MIGR1 or p210MINVneo viruses. Where indicated, *Ptpn11* deletion was induced with poly I:C (125µg x 2 times) for *in vivo* experiments or 4-OH (1µM) for *in vitro* studies. Transduced cells were isolated by FACS (for p210MIGFP) or G418 selection (for p210MINVneo), starved for 2 hours in IMDM-2%FBS (myeloid cells) or OptiMEM-2%FBS (lymphoid cells), and lysed in RIPA buffer.^47^ Lysates were resolved by SDS-PAGE, and immunoblotting was performed as described.^21^ Anti-phospho-STAT5 (Tyr694), -phospho-CRKL (Tyr207), -phospho-(Ser473) and -total ART, -phospho-(Thr202/Tyr204) and -total p44/42 ERK1/2, -phospho-S6 (S235/236), SRC, and -phospho-SRC (Y416), which cross-reacts with the other SFKs, were from Cell Signaling Technology. Anti-ABL1 and -SHP2 were from Santa Cruz Biotechnology. Antibodies were used at each manufacturer’s recommended concentrations.

### RNA-seq

RNA (150 ng) was reverse-transcribed using the Illumina TruSeq Stranded mRNA kit. cDNA libraries were size-indexed and validated using an Agilent Bioanalyzer, and concentrations were confirmed by qPCR. For the 16 samples, each of the 4 libraries (1 WT SHP2^-/-^, 1 WT SHP2′ 1 BCR-ABL1 SHP2^+/+^, and 1 BCR-ABL1 SHP2^-/-^) was loaded onto an Illumina cBot for cluster generation, and the flow cell was subjected to 100 cycles of paired-end sequencing on an Illumina HiSeq 2000. Alignment was performed with Bowtie, using default parameters. Gene expression levels were estimated using Cufflinks and subjected to quantile normalization.^48^ Batch effect adjustment was performed by Combat.^49^ Differentially expressed genes between samples were identified by voom-limma.^50^ Enrichment analyses for GO terms or transcription factor effectors were performed with DAVID or Enrichr, respectively.^51,52^ GSEA was implemented using software from the Broad Institute.

### Quantitative RT-PCR

Total RNA was prepared using the RNeasy minikit (Qiagen), and reverse-transcribed using SuperScriptIII (Invitrogen). SYBR Green-based gene expression analyses (BIO-RAD) were conducted according to the manufacturer’s instructions. Each sample was measured in triplicate, and relative expression was normalized to *Tbp.*

### Statistics

Survival curves were compared by the log-rank test. Signaling intensities in myeloid and lymphoid cells were compared by two-sided *t*-tests. Corrections for multiple comparisons were performed by using the Benjamini-Hochberg procedure. Sample size was determined to ensure at least 80% power to detect the difference of interest, based on the empirically estimated variance in each group.

### Study approval

All animal studies were approved by the University Health Network Animal Care Committee (Toronto, Ontario, Canada) and performed in accordance with the standards of the Canadian Council on Animal Care.

## Results

### SHP2 deficiency attenuates induction of CML-like MPN by BCR/ABL

BCR-ABL1 induces cytokine-independent myeloid progenitor colonies,^53^ and BCR-ABL1-transformed hematopoietic stem and progenitor cells (HSPCs), transplanted into irradiated syngeneic recipient mice, induce CML-like MPN.^13^ To address the role of SHP2 in BCR-ABL1-evoked myeloid disease, we transduced BM from *Ptpn11^fl/fl^* mice^41^ with p210MIGFP- or p210MIGFPCre-based retroviruses that co-express BCR-ABL1 with GFP or GFP-CRE, respectively. GFP-CRE induces floxed allele deletion, resulting in absence of SHP2. Lin^-^ BM cells from WT or *Ptpn11^fl/fl^* mice were infected with parental pMIGR1, p210MIGFP, or p210MIGFPCre virus, GFP^+^ cells were isolated by FACS and seeded in methylcellulose-based media, and cytokine-independent colonies were counted 1 week later. As expected, p210MIGFP-infected WT and *Ptpn11*^*fl/fl*^ BM cells yielded similar numbers of cytokine-independent myeloid colonies. By contrast, myeloid colony formation by p210MIGFPCre-infected *Ptpn11*^*fl/fl*^ BM cells was impaired markedly, indicating that SHP2 is required for BCR-ABL1-induced myeloid transformation (Figure 1a). Notably, p210MIGFPCre-infected WT BM yielded fewer cytokine-independent colonies than did p210MIGFP-infected BM, suggesting that constitutive CRE expression might impair myeloid transformation, consistent with previous reports of CRE toxicity.^54^

**Figure 1.**
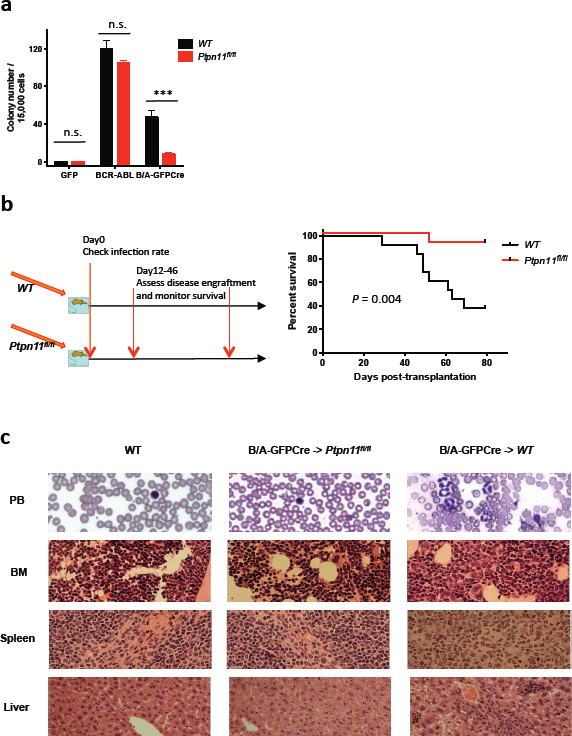
SHP2 is required for BCR-ABL1-evoked myeloid transformation and initiation of CML-like MPN. (a) Cytokine-independent colony formation of *WT* or *Ptpn11*^*fl/fl*^ cells infected with pMIGR1 (GFP), p210MIGFP (BCR-ABL), or p210MIGFPCre (BCR-ABL-GFPCre) virus (n = 4). (n.s.=not significant; *** *P <* 0.001). *WT* cells infected with p210MIGFP yielded more cytokine-independent colonies than infected with p210MIGFPCre *(P <* 0.001). (b) Scheme of experiment to assess the requirement for SHP2 in initiation of CML-like MPN (left), and Kaplan-Meier curve of recipients of *WT* or *Ptpn11*^*fl/fl*^ donor cells infected with p210MIGFPCre virus (right, n = 13). Recipients of the *Ptpn11*^*fl/fl*^ donor cells survived significantly longer *(P =* 0.004; log-rank test). A similar trend showing that p210GFPCre recipients survive longer than p210GFP recipients was reported by Walz *et al..^43^* (c) Histology of peripheral blood, bone marrow, spleen, and liver in typical WT mouse or recipients of *WT* or *Ptpn11*^*fl/fl*^ cells infected with p210MIGFPCre virus (400x amplification).

We next transplanted p210MIGFP-Cre transduced, Lin^-^ BM cells from WT or *Ptpn11*^*fl/fl*^ mice into sub-lethally irradiated WT recipients (Figure 1b). Recipients of BCR-ABL1-expressing WT BM cells developed fatal, CML-like MPN, characterized by leukocytosis with maturing myeloid cells, splenomegaly, and leukemic cell infiltrates in spleen and liver (Figure 1b,c and Supplementary Figure S1). Disease latency and histopathology were comparable to that reported previously.^43^ By contrast, the *Ptpn11*^*fl/fl*^ cohort had markedly extended survival, reduced leukocytosis and splenomegaly, and diminished leukemic cell infiltration (Figure 1b,c and Supplementary Figure S1). Notably, 12 days post-transplant, peripheral blood and spleen leukemic cell number were much higher in recipients of WT BM cells, compared with *Ptpn11*^*fl/fl*^ BM recipients (Supplementary Figure S1a,b), while BM engraftment was comparable (Supplementary Figure S1c). Therefore, SHP2 is required for initiation of CML, rather than homing to, or establishment in, the BM niche.

### SHP2 is required for maintenance of BCR-ABL1-evoked CML-like MPN

To ask if SHP2 is required for CML maintenance, we crossed *Ptpn11*^*fl/fl*^ mice to mice expressing *Cre* under the control of the type I interferon-inducible *Mx* promoter (*MxCre*), and infected BM cells from *MxCre* or *MxCre;Ptpn11^fl/fl^* donors with p210MIGFP virus. Between 11-15 days following transplantation of infected BM cells into WT recipients, two thirds of the mice in each cohort were injected with poly I:C to induce *Cre* expression in donor cells (Figure 2a). All recipients of *MxCre* BM or *MxCre;Ptpn11^fl/fl^* BM that had not been poly I:C-treated, succumbed to CML-like MPN within 30 days post-BM transplantation (BMT). Poly I:C-treated recipients of *MxCre;Ptpn11^fl/fl^* BM cells exhibited delayed/diminished disease and longer survival (Figure 2a). Leukemic cell levels in peripheral blood were reduced substantially (Figure 2a and Supplementary Figure S2a), and WBC counts were normalized (Supplementary Figure S2b). All recipients that developed CML-like MPN despite poly I:C induction had incomplete *Ptpn11* deletion in donor leukemic cells (Supplementary Figure S3). Hence, SHP2 is also necessary for CML-like MPN maintenance.

**Figure 2.**
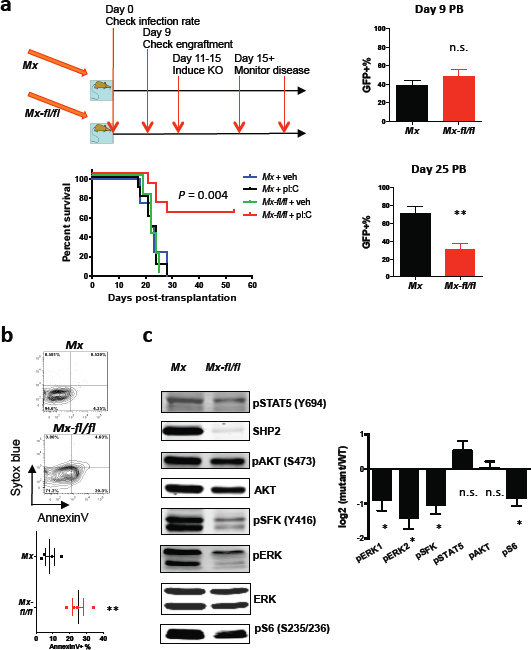
SHP2 is required for CML maintenance. (a) Left: Scheme of experiment and Kaplan-Meier curve for recipients of *MxCre* (*Mx*) or *MxCre;Ptpn11^fl/fl^* (*Mx-fl/fl)* bone marrow infected with p210MIGFP virus, with (n = 10) or without (n = 5) poly I:C induction. The *Mx-fl/fl* + poly I:C group survived significantly longer than the other groups *(P* = 0.004; log-rank test). Right: Peripheral blood leukemic engraftment on day 9 and day 25 post-transplantation in *Mx +* pI:C and *Mx-fl/fl* + pI:C groups. Note that engraftment is comparable before poly I:C induction (day 9), but different after *Ptpn11* deletion (day 25, *P* < 0.01). (b) GFP^+^Lin^-^Scal^+^cKit^+^ splenocytes were isolated from poly I:C-induced recipients of *Mx* or *Mx-fl/fl* donor cells, and stained with Annexin V and Sytox blue (n = 4). Note that apoptosis is significantly higher in *Mx-fl/fl* than *Mx* group *(P <* 0.01). (c) Left: Representative immunoblot of poly I:C-induced *Mx* or *Mx-fl/fl* BCR-ABL1^+^ myeloid cells. Similar results were obtained in ≥5 biological replicates. Right: Statistical analysis of ratio of phosphorylated to total signaling protein levels from ≥5 biological replicates of the experiment shown in G *(*P <* 0.05, n.s., not significant; 2-sided Student *t* tests).

SHP2 is required for the survival of WT primitive hematopoietic cells,^31^ so we assessed its role in primitive CML cells. GFP^+^Lin^-^Scal^+^cKit^+^ splenocytes from poly I:C-treated recipient mice were stained with Sytox blue and Annexin V, and analyzed by flow cytometry. Apoptosis was increased markedly in SHP2-deficient, compared with control, cells (Figure 2b).

To study the role of SHP2 in BCR-ABL1-evoked myeloid signaling, we infected SHP2-expressing or -deficient Lin^-^ BM cells with p210MIGFP retrovirus, isolated infected cells, and performed immunoblotting. In the absence of SHP2, phospho-ERK1 and -ERK2 levels were reduced (Figure 2c), as was phospho-S6-S235/236; the latter sites can be phosphorylated by the ERK-dependent kinase RSK2, as well as by mTORC1.^55^ These results are consistent with the requirement for RAS-ERK signaling in BCR-ABL1-induced CML.^56–58^ In addition, phospho-SFK-Y416, which marks the catalytically activated form of SFKs, was reduced in the absence of SHP2. By contrast, we did not see significant changes in phospho-AKT or phospho-STAT5 levels.

### SHP2 is required for Ph^+^ B-ALL

As expected, BCR-ABL1-expressing, IL-7-stimulated bulk BM cells evoked B-ALL in lethally irradiated syngeneic recipients.^12^ To probe the role of SHP2 in Ph^+^ B-ALL initiation, we transplanted IL-7-stimulated BM from *WT* or *Ptpn11^fl/fl^* donors transduced with p210MIGFPCre virus into lethally irradiated syngeneic recipients. Compared with recipients of BCR-ABL1-infected WT BM, recipients of transduced *Ptpn11*^*fl/fl*^ BM had much lower levels of BCR-ABL1^+^ B lineage engraftment in their peripheral blood (Figure 3a) and survived longer (Figure 3b). Hence, SHP2 is required for effective initiation of BCR-ABL1^+^ B-ALL. Consistent with the effects of GFP-CRE in myeloid cells, we observed less severe disease in recipients of p210MIGFPCre, compared with recipients of p210MIGFP (Figure 3a), BM, possibly reflecting CRE toxicity.

**Figure 3.**
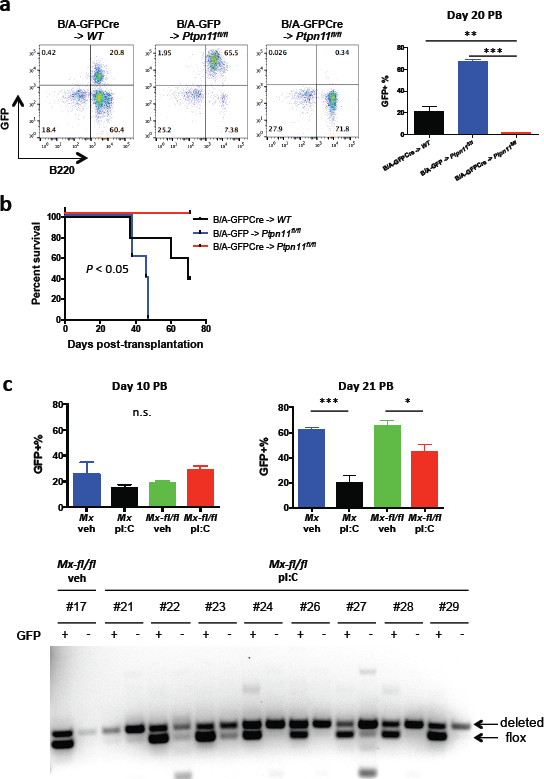
SHP2 is required for initiation, and most likely, for maintenance, of Ph^+^ B-ALL. (a) Leukemic engraftment in recipients of *WT* or *Ptpn11^fl/fl^* donor cells infected with p210MIGFP (B/A-GFP) or p210MIGFPCre (B/A-GFPCre) retrovirus, assessed on day 20 post-transplantation. Engraftment is significantly lower in B/A-GFPCre -> *Ptpn11*^*fl/fl*^ than in the other two groups (n = 5; mean ± s.e.m.; *\*\*P* < 0.01, *\*\*\*P <* 0.001; *N=* 5 each; one-way ANOVA with Bonferroni post-test). (b) Kaplan-Meier curve of recipients of *WT* or *Ptpn11*^*fl/fl*^ donor cells infected with p210MIGFP (B/A-GFP) or p210MIGFPCre (B/A-GFPCre) retrovirus (n = 5). Survival was longer in B/A-GFPCre -> *Ptpnll^fl/fl^* mice *(P <* 0.05; log-rank test). (c) Upper: Peripheral blood leukemic engraftment on day 10 and day 21 post-transplantation in *Mx* and *Mx-fl/fl* groups (n = 5 for vehicle control groups and n = 10 for poly I:C groups; mean ± s.e.m.). Note that engraftment was comparable before, but significantly lower after poly I:C treatment *(*P <* 0.05, *\*\*\*P <* 0.001; one-way ANOVA with Bonferroni post-test), indicating the effect of poly I:C treatment itself and complicating the interpretation of the essentiality of SHP2 in disease maintenance. Lower: GFP^+^ (leukemic) or GFP^-^ (WT) B-lymphoid cells were isolated from peripheral blood of recipients of *Mx-fl/fl* donor cells, treated with vehicle or poly I:C, as indicated. *Ptpn11* deletion was assessed by PCR of genomic DNA. Deleted and *floxed* alleles are as indicated. Bands that are not “deleted” or “*floxed*” are likely non-specific. Note that recipients in the *Mx-fl/fl* + pI:C group exhibited more efficient deletion in GFP’ than GFP^+^ cells.

We next attempted to assess the effect of *Ptpn11* deletion on B-ALL maintenance. Bulk BM cells from *MxCre* or *MxCre;Ptpn11^fl/fl^* donors were infected with p210MIGFP virus, and transplanted into lethally irradiated, syngeneic recipients. After B-ALL developed, half of the recipients in each cohort were treated with poly I:C to induce *Ptpn11* deletion. Unexpectedly, poly I:C alone significantly alleviated disease (Figure 3c), consistent with an interferon-induced inhibitory effect against B-ALL. These effects prevented assessment of the role of SHP2 on established B-ALL. We also generated mice (*Rosa26-CreER;Ptpn11^fl/fl^*) expressing a CRE-ER fusion protein, and used tamoxifen to induce *Ptpn11* deletion. Unfortunately, the floxed allele did not delete efficiently, and although tamoxifen-treated mice showed a small increase in survival, *Ptpn11*-undeleted BCR-ABL blasts rapidly outgrew (data not shown).

Instead, we compared the abundance of *Ptpn11*-replete and -deleted cells in the same mouse: if SHP2 is required for disease maintenance, then *Ptpn11*-replete leukemic cells should out-compete *Ptpn11*-deficient cells. *Ptpn11* deletion was assessed by PCR of GFP^+^ (BCR-ABL1^+^) and GFP^-^ (uninfected) peripheral blood cell DNA from poly I:C-induced *MxCre;Ptpn11^fl/fl^* recipients. *Ptpn11* deletion was much more efficient in GFP^-^, than in GFP^+^, cells (Figure 3c), indicating that SHP2 is required specifically for BCR-ABL1^+^ B-lymphoid proliferation/survival.

### SHP2 deficiency selectively affects cycling of BCR-ABL1^+^ pre-B cells

We compared the effects of SHP2 deficiency on WT and BCR-ABL1^+^ pre-B cell proliferation *in vitro. Rosa26-CreER;Ptpn11^fl/fl^* pre-B cells were cultured in IL7-supplemented OptiMEM-10%FBS, and *Ptpn11* deletion was induced with 4-OH in half of the cells. Unexpectedly, SHP2-deficient WT cells proliferated at the same rate as untreated controls (Figure 4a); hence, SHP2 is dispensable for WT pre-B cell proliferation *in vitro.* Furthermore, *Cd19^tm1(cre)Cgn^;Ptpn11^fl/fl^* (Supplementary Figure S4) mice exhibited grossly normal B cell development and agonist responses, and preliminary studies of *Cd79a^tm1(cre)Reth^;Ptpn11^fl/fl^* mice yielded similar results. Next, we cultured BCR-ABL1^+^ *Rosa26-CreER* or BCR-ABL1^+^ *Rosa26-CreER;Ptpn11^fl/fl^* pre-B cells, and induced *Ptpn11* deletion in the latter with 4-OH. Remarkably, SHP2-deficiency selectively abrogated the proliferation of BCR-ABL1^+^ pre-B cells (Figure 4a and Supplementary Figure S5). The allosteric SHP2 inhibitor SHP099^59^ also markedly suppressed BCR-ABL1^+^ cell proliferation (Supplementary Figure S6). Hence, SHP2 is required for BCR-ABL1^+^-evoked, but not WT, pre-B cell proliferation. SHP2 deficiency did not increase apoptosis in BCR-ABL1^+^ cells (Figure 4b), but instead, caused increased G0 and decreased G1 and S-G2-M cells (Figure 4c).

**Figure 4.**
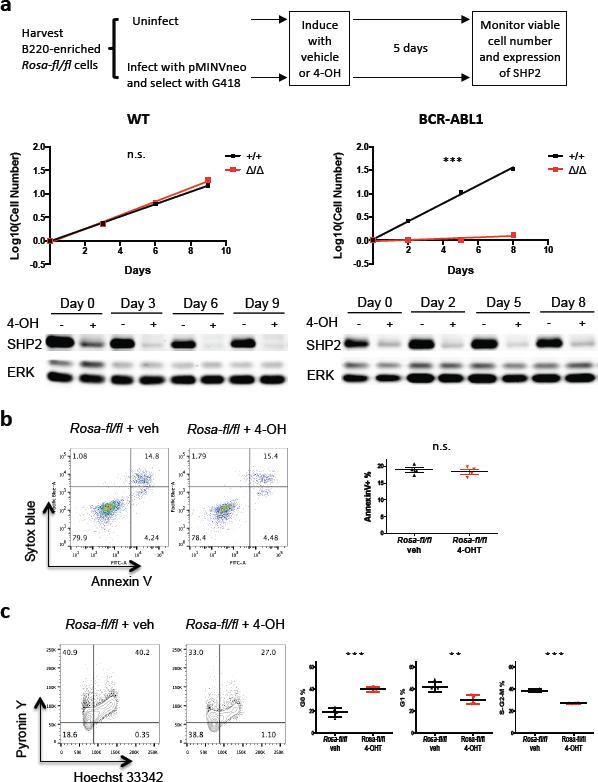
SHP2 is required for the proliferation of BCR/ABL1^+^ pre-B cells *in vitro*. (a) Bone marrow cells were harvested from *Rosa26-CreER;Ptpn11^fl/fl^* mice, and B-lineage cells were enriched as in Methods. Cells were cultured in the presence of IL7, left uninfected or infected with p210MINVneo, and then selected with G418. Two days post-infection, cells in each cohort were treated with vehicle or 4-OH to induce *Ptpn11* deletion. Viable cell number and SHP2 levels were monitored starting on day 5 post-deletion (“Day 0” in growth curves and immunoblots). WT *Rosa26-CreER;Ptpn11^fl/fl^* pre-B cells with intact (+/+) or deleted (Δ/Δ) *Ptpn11* were culture and cell number was monitored (n = 4). Proliferation rates were not significantly different between +/+ and Δ/Δ cells. BCR-ABL1^+^ *Rosa26-CreER;Ptpnll^fl/fl^* pre-B cells with intact (+/+) or deleted (Δ/Δ) *Ptpn11* were cultured, and cell number was monitored (n = 4). Note that the proliferation of SHP2-deficient BCR-ABL1^+^ pre-B cells was impaired *(***P <* 0.001; *F* test). Efficiency of *Ptpn11* deletion was similar between WT and BCR-ABL1^+^ cells, (b) BCR-ABL1^+^ *Rosa26-CreER;Ptpn11^fl/fl^* pre-B cells treated with vehicle or 4-OH were analyzed by flow cytometry for Annexin V and Sytox blue (n = 4; mean ± s.d.; n.s. = not significant; two-sided t test). (c) BCR-ABL1^+^ *Rosa26-CreER;Ptpnl* pre-B cells treated with vehicle or 4-OH were stained with Hoechst 33342 and Pyronin Y (n = 4; mean ± s.d.; *\*\*P <* 0.01,***P < 0.001; two-sided t test with Benjamini-Hochberg adjustment for multiple comparisons.)

### SFK and MEK-ERK signaling contribute to the differential SHP2 requirement

We next assessed the effects of SHP2 on WT and BCR-ABL1^+^ pre-B cell signaling. In WT cells, SHP2 deficiency reduced phospho-ERK1 and phospho-ERK2 levels (Figure 5a), indicating that WT pre-B cells are insensitive to a reduction of ERK signaling of this magnitude. In WT pre-B cells, phospho-AKT (S473), phospho-STAT5 (Y694), and phospho-SFK (Y416) levels were not affected by SHP2 deficiency. In BCR-ABL1^+^ pre-B cells, *Ptpn11* deletion also led to significantly lower levels of phospho-ERK1, phospho-ERK2, but comparable levels of phospho-AKT (S473) and phospho-STAT5 (Y694) (Figure 5a). However, *Ptpn11* deletion also led to compromised SFK activation, as judged by phospho-SFK (Y416), only in BCR-ABL1^+^ pre-B cells (Figure 5a).

**Figure 5.**
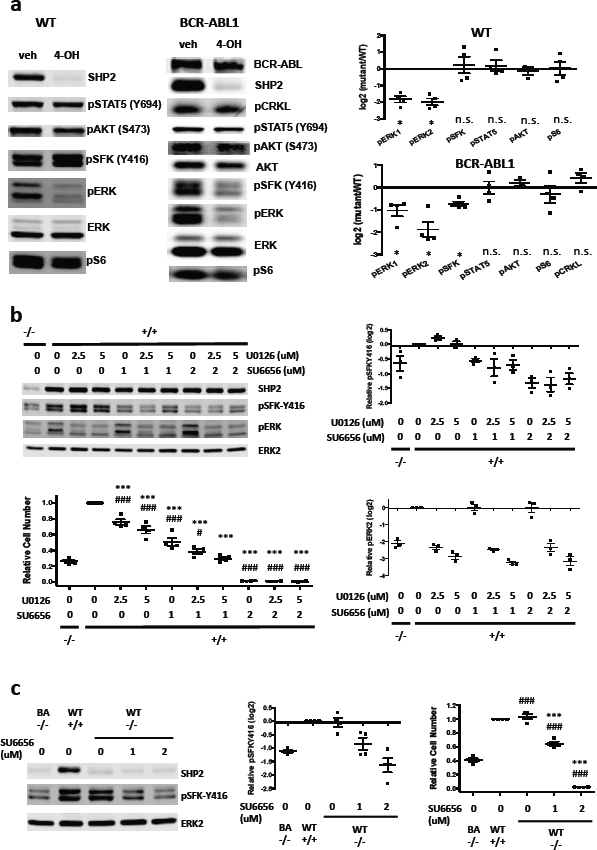
SFKs and ERK mediate the SHP2 requirement in BCR-ABL1^+^ pre-B cells. (a) Signaling in WT or BCR-ABL1^+^ *Rosa26-CreER;Ptpn11^fl/fl^* pre-B cells treated with vehicle or 4-OH. Similar results were obtained in 4 biological replicates. (mean ± s.e.m.; *\*P <* 0.05; 2-sided *t* test with Benjamini-Hochberg adjustment for multiple comparisons). (b) SHP2-replete BCR-ABL1^+^ pre-B cells were treated with the MEK/ERK inhibitor (JO 126 and/or the SFK inhibitor SU6656 at different concentrations, and compared with SHP2-deficient BCR-ABL1^+^ pre-B cells on signaling (1-hour drug treatment, n = 3) or viable cell number (3-day treatment, n = 4). (JO 126 specifically affects phospho-ERK, and SU6656 specifically affects phospho-SFK. Viable cell numbers post-drug treatment were compared with SHP2-replete BCR-ABL1^+^ vehicle control (denoted by “*”) or SHP2-deficient control (denoted by “#”) groups. The significance of the cell number difference between combination treatment and SHP2 deficiency is decreased compared with the difference between single-agent treatment and SHP2 deficiency. (mean ± s.e.m.; *P <* 0.05 between U0126_(2.5µM)_ + SU6656n_(1µm)_) vs SHP2^-/-^, and the difference is not significant between U0126_(5µM)_ + SU6656_(1µM)_ vs SHP2^-/-^) (C) SHP2-deficient WT pre-B cells were treated with SU6656 (0, 1, 2 µM), and compared to SHP2-replete WT pre-B cells or SHP2-deficient BCR-ABL1^+^ pre-B cells on signaling (1-hour treatment) or viable cell number (3-day treatment). Total ERK2 serves as a loading control. Viable cell numbers from SHP2-deficient WT groups were compared with SHP2-replete WT (denoted by “*”) or SHP2-deficient BCR-ABL1^+^ (denoted by “#”) groups (n = 4). 1 µM SU6656 treatment recapitulates most of the proliferation inhibition by SHP2 deficiency, although the difference is still significant (0.65±0.06 vs 0.41±0.05, mean ± s.e.m.). (*\*\*\*P <* 0.001, *#P <* 0.05, *###P* < 0.001; ANOVA with Bonferroni post-test multiple comparison)

To ask if decreased MEK/ERK and/or SFK activity contribute to the requirement for SHP2 for BCR-ABL1^+^ pre-B cell proliferation, we treated (SHP2-replete) BCR-ABL1^+^ pre-B cells with MEK/ERK (U0126) and/or SFK-selective (SU6656)^60^ inhibitors. At 1µM, SU6656 inhibited SFK activity (as assessed by phospho-SFK Y416) to levels similar that caused by SHP2 deficiency (Figure 5b). This dose also impaired BCR-ABL1^+^ pre-B cell proliferation, although not by as much as *Ptpn11* deletion (Figure 5b). At 2.5µM, U0126 diminished phospho-ERK2 to levels comparable to those in SHP2-deficient cells, and had a smaller, but significant, effect on proliferation (Figure 5b). Combined SU6656-U0126 treatment recapitulated the proliferation inhibition caused by SHP2 deficiency. Treatment with another selective SFK inhibitor, PP2, yielded similar results (data not shown). Therefore, decreased proliferation of SHP2-deficient, BCR-ABL1^+^ pre-B cells results from the combined effects of impaired SFK and ERK activation, with the former playing a more important role. SU6656 did not affect ERK activation and U0126 did not affect SFK activation, so these pathways function independently downstream of SHP2 in pre-B cells.

The ability of WT pre-B cells to activate SFKs without SHP2 might explain why they do not require SHP2 to proliferate. Treatment of SHP2-deficient (BCR-ABL1-negative) pre-B cells with SU6656 (1µM) reduced phospho-SFK to levels slightly more than did SHP2-deficiency in BCR-ABL1^+^ cells, whereas 2µM SU6656 lowered p-SFK levels to a slightly greater extent. Remarkably, these doses of SU6656 inhibited SHP2-deficient WT pre-B cell proliferation to an extent slightly less than, or slightly more than, did SHP2-deficiency in BCR-ABL1^+^ pre-B cells (Figure 5c). Persistent SFK signaling probably explains why SHP2 is dispensable for WT pre-B cell proliferation.

### Structural determinants of SHP2 required for BCR-ABL1^+^ pre-B cell proliferation

To identify sub-domains of SHP2 required for BCR-ABL1^+^ signaling in pre-B cells, we compared proliferation and signaling in SHP2-sufficient (veh) or -deficient BCR-ABL1^+^ pre-B cells reconstituted with empty vector (4-OH), WT SHP2 (WT), the phosphatase-inactive C459E mutant (CE), or a mutant lacking both C-terminal tyrosine residues, Y542F/Y580F (Y2F). WT SHP2 rescued proliferation and ERK and SFK activity. By contrast, the CE or Y2F mutant did not restore proliferation or signaling (Figure 6), even though each was expressed at levels similar to WT SHP2. SH2 domain engagement is also essential, given the effects of SHP099 (Supplementary Figure S6), which locks SHP2 in the closed form and prevents N-SH2/p-Tyr peptide binding.^59,61^

**Figure 6.**
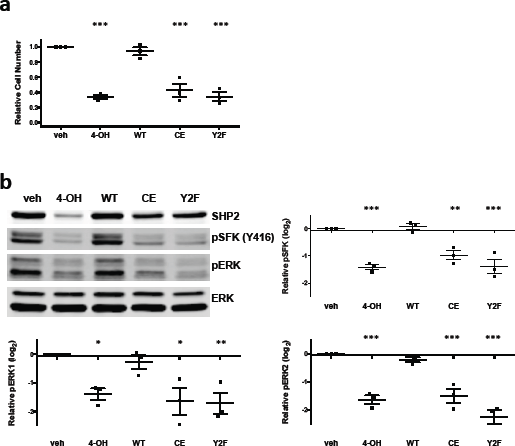
SHP2 domains required for signaling and proliferation. (a) Relative viable cell number after 2-day cultures, pooled from 3 biological replicates. The proliferation of *Rosa26-CreER;Ptpnl*BCR-ABL1^+^ cells subjected to the indicated treatments was assessed: (1) empty vector infection followed by vehicle treatment (veh), (2) empty vector infection followed by 4-OH treatment (4-OH), (3-5) reconstitution with SHP2 variants (WT, C459E or Y542F/Y580F) followed by 4-OH treatment. Viable cell numbers were compared with the vehicle group, (b) Comparison of phospho-SFK(Y416) and phospho-ERK in *Rosa26-CreER;Ptpn11^fl/fl^* BCR-ABL1^+^ cells from experiments in panel (a). Note that reconstitution with WT SHP2, but not the C459E or Y542F/Y580F mutant, restores ERK and SFK activity. (n = 3; mean ± s.e.m.; *\*P <* 0.05, *\*\*P <* 0.01, *\*\*\*P <* 0.001; ANOVA with Bonferroni post-test)

### Distinct effects of SHP2 on transcriptional profiles of BCR-ABL1^+^ and WT pre-B cells

To gain further insight into the role of SHP2 in WT and BCR-ABL1^+^ pre-B cells, we performed RNA-seq. Unsupervised hierarchical clustering and principal component analysis (PCA) revealed good separation between both WT and BCR-ABL1^+^ cells and between SHP2-deficient and -replete cells, respectively (Supplementary Figure S7). As expected ^62,63^, BCR-ABL1^+^ cells (compared with WT) showed up-regulation of ERK targets and down-regulation of interferon-alpha response genes (Supplementary Figure S8 and Supplementary Tables S1-S2). Supervised comparisons revealed that SHP2 significantly regulates the expression of ~770 genes in WT, and >1900 genes in BCR-ABL1^+^, pre-B cells (Figure 7a). Also as expected, SHP2 deficiency decreased ERK-regulated gene expression in WT and BCR-ABL1^+^ samples (Supplementary Table S3). TLR-regulated genes also were decreased by SHP2-deficiency in both contexts. Shared up-regulated genes were not enriched for any curated Reactome pathway (Supplementary Table S4).

**Figure 7.**
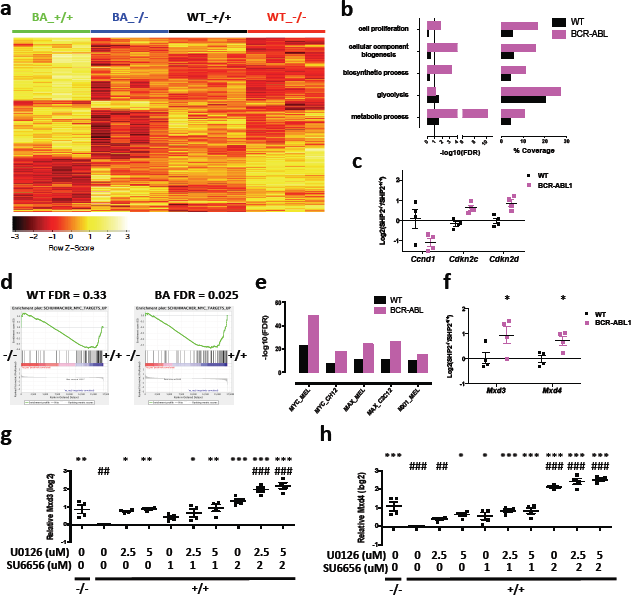
SHP2 affects distinct transcriptional programs in WT and BCR-ABL1^+^ pre-B cells. (a) Hierarchical clustering of RNA-seq data from: WT SHP2^+/+^ (WT_+/+), WT SHP2′ ‘ (WT -/-), BCR-ABL SHP2^+/+^ (BA_+/+), and BCR-ABL SHP2“” (BA -/-), based on genes significantly regulated by SHP2 in either WT or BCR-ABL1^+^ cells (4 samples per group). (b) SHP2-regulated genes in WT or BCR-ABL1^+^ cells enrich for different GO terms. Bar graph (left) denotes significance of enrichment for each GO term in the SHP2-regulated gene sets. Bar graph on the right denotes the percentage of SHP2-regulated genes in each GO category in WT or BCR-ABL1^+^ cells. (c) Validation of differential expression of genes involved in cell cycle regulation by q-RT-PCR (n = 4). *Ccnd1*, *Cdkn2c*, and *Cdkn2d* are regulated by SHP2 in in BCR-ABL1^+^, but not WT, pre-B cells. (d) GSEA plots of MYC-induced signatures from WT_-/-versus WT_+/+ or BA_-/-versus BA_+/+ cells. (e) Binding sites for MYC and its interactors are enriched differentially in genes down-regulated by SHP2 deficiency in WT or BCR-ABL1^+^ pre-B cells. (f) qRT-PCR validation of genes involved in regulation of MYC transcription (n = 4). The log ratios of *Mxd3* or *Mxd4* transcription levels in SHP2-deficient vs SHP2-replete cells are shown. *Mxd3* and *Mxd4* mRNA levels are not regulated by SHP2 deficiency in WT pre-B cells, but are significantly up-regulated in BCR-ABL1^+^ pre-B cells. (g-h). BCR-ABL1^+^ SHP2-deficient (-/-) or replete (+/+) pre-B cells were treated with U0126 and/or SU6656 at indicated concentrations for 2 days. Relative mRNA levels for *Mxd3* (g) or *Mxd4* (h) were quantified (n = 4). Transcription levels were were compared with SHP2-replete BCR-ABL1^+^ vehicle control (denoted by “*”) or SHP2-deficient control (denoted by “#”) groups. *(*P <* 0.05, *\*\*P <* 0.01, *\*\*\*P* < 0.001, *##P <* 0.01, *###P* < 0.001; ANOVA with Bonferroni post-test).

Next, we compared BCR-ABL1^+^ SHP2-deficient versus SHP2-expressing cells and WT SHP2-deficient versus SHP2-expressing cells, respectively. SHP2 deficiency in normal pre-B cells affected genes annotated by the terms metabolic process and glycolysis (Figure 7b and Supplementary Table S5). In BCR-ABL1^+^ cells, SHP2-deficiency also affected genes for metabolic and biosynthetic processes; in addition, cell-proliferation genes were altered (Figure 7b,c and Supplementary Table S6). We identified, and validated by qRT-PCR, differential effects of SHP2 deficiency on cell cycle regulators in WT and BCR-ABL1^+^ cells. *Ccnd1* (encoding CYCLIN D1) expression was decreased, while genes encoding the CDK inhibitors p18ARF and p19ARF (*Cdkn2c* and *Cdkn2d*) were increased preferentially by SHP2 deficiency in BCR-ABL1^+^ cells (Figure 7c). These results comport with the cell cycle impairment seen in SHP2-deficient BCR-ABL1^+^ pre-B cells (Figure 4c).

Consistent with GAB2 regulating multiple signal relay molecules,^15,64^ more genes and gene sets were affected by GAB2 ^21^, than by SHP2, deficiency (Supplementary Figure S9a and Supplementary Tables S7-S8). Nevertheless, the top gene sets affected by either deficiency showed strong overlap and similar FDR q-values (Supplementary Figure S9b). Gene sets down-regulated only by GAB2 deficiency included multiple DNA repair/synthesis-related and metastasis-related gene sets, among others (Supplementary Tables S7-S8).

GSEA revealed down-regulation of MYC-induced signatures in SHP2- (and GAB2-) deficient, BCR-ABL1^+^ cells, but no significant alteration in SHP2-deficient WT cells (Figure 7d, Supplementary Tables S7 & S9). Enrichr analysis showed that genes down-regulated by SHP2 deficiency in BCR-ABL1^+^ cells were highly enriched for binding sites for MYC and its cofactors (Supplementary Table S12). Genes down-regulated in SHP2 deficient WT cells also were enriched for MYC/co-factor binding sites, but with much lower overlap and significance (Figure 7e and Supplementary Table S13). Intriguingly, multiple MYC targets were regulated in opposite directions in SHP2-deficient BCR-ABL1^+^ and WT pre-B cells. For example, glycolytic gene expression was increased in WT cells, but down-regulated in BCR-ABL1^+^ cells (Supplementary Figure S10). Hence, SHP2 deficiency has quantitatively and qualitatively distinct effects on MYC-dependent transcriptome (“MYC regulome”) in BCR-ABL1^+^ (compared with WT) pre-B cells.

Gene sets up-regulated by SHP2 deficiency in BCR-ABL1^+^ and WT cells differed widely. Interferon response genes were among the top up-regulated in BCR-ABL1^+^ cells (Supplementary Figure S11 and Supplementary Table S10), whereas hypoxia response genes were the top up-regulated sets in WT cells (Supplementary Table S11). Up-regulated genes in WT and BCR-ABL1 contexts were enriched comparably for binding sites for EP300, ETS1, and several other transcription factors/regulators (Supplementary Tables S14-S15). Genes with binding sites for CHD2 and ZKSCAN1 were enriched preferentially in BCR-ABL1+ cells, whereas those with RAD21 binding sites were enriched preferentially in WT cells (Supplementary Tables S14-S15).

Given the key role of MYC in cell proliferation,^65^ the selectively impaired proliferation of SHP2-deficient BCR-ABL1+ pre-B cells probably reflects alteration of the MYC regulome, at least in part. We asked why MYC target genes are regulated differentially in SHP2-deficient WT and BCR-ABL1^+^ cells. *Myc* mRNA levels were unaffected by SHP2-deficiency in WT or BCR-ABL1^+^ cells (Supplementary Figure S12a). MYC stability is regulated by ERK^66^ and, consistent with impaired ERK activation in SHP2-deficient WT and BCR-ABL1^+^ pre-B cells, MYC levels were decreased in both, but to similar extents (Supplementary Figure S12b). By contrast, *Mxd3* and *Mxd4* were up-regulated by SHP2 deficiency only in BCR-ABL1^+^, but not in WT cells (Figure 7f and Supplementary Figure S13). These increases were recapitulated by co-inhibition of ERK and SFKs in SHP2-replete BCR-ABL1+ cells (Figure 7g,h). MXD3 and MXD4 compete with MYC for binding to MAX and inhibit transcription of a subset of MYC targets involved in proliferation and transformation.^67^ Failure to repress *Mxd3* and *Mxd4* transcription might explain the more profound impairment of MYC-dependent transcription in SHP2-deficient BCR-ABL1 pre-B cells.

## Discussion

Our genetic approach enabled us to assess the role of SHP2 in BCR-ABL1-induced neoplasia in well-characterized mouse models. We find that SHP2 is essential for initiation and maintenance of CML-like MPN, and initiation and, most likely, maintenance, of Ph^+^ B-ALL. Similar to its effects in WT HSCs,^31,32^ SHP2 is required for the survival of phenotypic CML-SCs. Intriguingly, SHP2, and in particular, SHP2 catalytic activity and its C-terminal tyrosine residues, is essential for BCR-ABL1^+^, but not WT, pre-B cell proliferation. SHP2 mediates ERK activation in WT and BCR-ABL+ cells, but its differential effect on proliferation correlates with, and is likely caused by, distinct effects on SFKs. Transcriptional profiling suggests that the combined effects of ERK and SFKs, perhaps by repressing *Mxd3/4* transcription, are needed to fully activate the MYC regulome.

Previous studies implicated SHP2 in critical events downstream of BCR-ABL1, but none addressed its role directly in a pathologically relevant context.^21,40^ For example, *PTPN11* knockdown reduces cytokine-independent colony formation by CD34^+^ CML cells *in vitro*,^68^ but its effects on CML and Ph+ B-ALL development remained unclear. SHP2 is required for BCR-ABL transformation of a yolk sac cell line,^40^ but BCR-ABL1-induced CML and B-ALL are typically adult diseases, and adult and embryonic hematopoiesis differ.^69^ GAB2, and its SHP2 binding sites, is essential for induction of Ph^+^ hematopoietic neoplasia in mice.^21^ Yet other signaling molecules might bind the same sites, and SHP2 uses other adaptors/interactors besides GAB2; hence, loss of GAB2/SHP2 binding might have underestimated the role of SHP2 in BCR-ABL1 disease.^24,25^ Notably, SHP2 binding sites on GAB2 are required for full activation of STAT5 and AKT,^21^ but SHP2 deficiency did not affect STAT5 or AKT activation in the current study. One potential explanation for this apparent discrepancy is that the SHP2 binding sites of GAB2 might be shared by other proteins. For example, SHP2 and SOCS3 both interact with pY759 of gp130.^70,71^ Likewise, other proteins might share the SHP2 binding sites of GAB2. Alternatively, whereas GAB2-bound SHP2 can activate AKT and STAT5, other signaling proteins might inhibit AKT and/or STAT5 via SHP2, such that the net effect of SHP2 depletion differs from disruption of GAB2-SHP2 interaction.

The relationship between SHP2 and the RAS-ERK pathway is well documented.^24,25^ In hematopoietic cells, gain-of-function *PTPN11* mutants induce RAS-ERK hyper-activation and can cause JMML. Conversely, *Ptpn11* deletion impairs ERK activation by growth factors and cytokines in HSCs and progenitors.^31,32^ SHP2 also is required for ERK activation in WT and BCR-ABL1 transformed myeloid and pre-B cells. Surprisingly, however, WT pre-B cells, unlike myeloid progenitors^31,32^ and multiple other proliferating cells,^72,73^ are agnostic to defective ERK activation. How pre-B cells avoid the requirement for ERK activation seen in almost all other proliferative cells and tissues awaits further investigation.

Regulation of SFKs by SHP2 also has been reported.^41,74,75^ For as yet unclear reasons, SHP2 is required for SFK activation in BCR-ABL1-transformed, but not WT pre-B cells. SHP2 also is required for BCR-ABL1^+^, but not WT, pre-B cell proliferation. Inhibitor studies suggest that SFKs (to a major extent) and ERK (to a lesser extent) contribute independently to this selective requirement. Likewise, SHP2-independent SFK activation in WT cells is likely a(the) major resason for the differential requirement for SHP2 in BCR-ABL1^+^ and WT pre-B cell proliferation. Consistent with our results, SFKs, although dispensable for CML, are essential for Ph^+^ B-ALL.^76^

Previous studies suggested that SHP2 catalytic activity is required for ERK and/or SFK activation.^41,77^ The C-terminal tyrosines (Y542 and Y580) modulate activation of ERK downstream of multiple RTKs,^78^ although their roles in SFK activation had not been defined. SH2 domain/pY peptide interaction also is required for myeloid transformation by leukemogenic SHP2 mutants.^79^ We found that SH2 domain engagement, PTP activity, and the C-terminal tyrosines are required for BCR-ABL1^+^ pre-B cell proliferation and full activation of ERK and SFKs by BCR-ABL1.

Our RNA-seq studies reveal transcriptional programs differentially regulated by SHP2 in WT and BCR-ABL1^+^ pre-B cells. GSEA and Enrichr strongly suggest that defective MYC-driven transcription is a major consequence of SHP2 deficiency and is affected differentially in BCR-ABL1^+^ and WT pre-B cells. Some MYC-regulated genes are affected by SHP2 deficiency in BCR-ABL1-transformed, but not WT B cells, whereas others, including glycolytic genes, are regulated in opposite directions. Failure to engage Warburg respiration, a common feature of malignant B cells and other tumor cells,^80,81^ might contribute to cell cycle arrest in SHP2-deficient BCR-ABL1+ pre-B cells. As MYC is a key transcriptional hub for B cell growth and proliferation control,^82^ differential MYC transcription is likely to be a(the) major reason for the impaired proliferation of BCR-ABL1^+^ SHP2-deficient B cells.

Down-regulated MYC activity also is likely mediated by decreased SFK and/or ERK activation. SFKs increase MYC mRNA/protein levels,^60,83–85^ and MYC is stabilized by ERK-mediated phosphorylation.^66,73,84^ Yet *Myc* upregulation/MYC stabilization does not explain the differential regulation of MYC-dependent transcription by SHP2. Instead, we find that SFKs and/or ERK regulate cofactors/regulators of MYC: SHP2 deficiency elevates *Mxd3* and *Mxd4* specifically in BCR-ABL1^+^ pre-B cells, and this regulation is recapitulated by co-inhibition of ERK and SFKs.

MXD3 and MXD4 can occupy the MYC E-box, inhibit MYC/MAX binding, and repress MYC-induced transformation.^86^ Serum or insulin induce phosphorylation-directed degradation of MXD1 in HeLa cells,^87^ but transcriptional regulation of MXDs has not been reported. Notably, inspection of RNA-seq data from PDGF-stimulated smooth muscle cells^88^ reveals that *MXD1, MXD3*, *MXD4*, and *MNT*, which encodes an MXD-like protein, are down-regulated. STK38 reportedly regulates MYC-dependent transcription in ST486 B cell lymphoma cells by affecting MYC turnover.^89^ In that dataset, however, *MXD4* and *MNT* also were up-regulated by *STK38* silencing. Re-analysis of other studies^90,91^ also reveals regulation of *MXDs* and/or *MNT* mRNA levels by various cytokines or growth factors. Hence, regulation of the *MXD* family^67^ might be a general, but under-appreciated, mechanism for modulating MYC-dependent transcription. MXD3/4 up-regulation by SHP2 deficiency in BCR-ABL1^+^ pre-B cells likely contributes to inhibition of MYC-mediated transcription.

Although altered MYC-dependent transcription probably is a major contributor to defective proliferation in SHP2-deficient BCR-ABL1 pre-B cells, Enrichr analysis suggests that several other transcription factors/regulators also are regulated differentially, including BHLHE40, ETS1, CHD1, and CHD2. Further studies are needed to determine whether these molecules contribute to the specific requirement of SHP2 in BCR-ABL1^+^ pre-B cells.

IFN-α responsive genes also were up-regulated by SHP2 deficiency specifically in BCR-ABL1^+^ cells. IFN-α has been used to treat Ph^+^ B-ALL (as well as CML) and improves survival.^92,93^ IFN-α also induces growth arrest, and sometimes differentiation, in human and murine B cell lines.^94–96^ We did not detect differential expression of drivers/markers of B cell differentiation, including *Ikzf1, Spi1*, *Pax5, Ebf1*, *Tcf3*, *Tnf13*, and *Tnf13b*^97–99^ (data not shown), so enhanced differentiation is unlikely to contribute to the effects of SHP2 deficiency in BCR-ABL1+ B-ALL.

In summary, we find that SHP2 is essential for the pathogenesis of BCR-ABL1-evoked myeloid and lymphoid neoplasia. SHP2 is required specifically for BCR-ABL1^+^, but not WT, pre-B cell proliferation, because it mediates SFK *and* ERK activation in BCR-ABL1^+^ pre-B cells, but only ERK activation in WT pre-B cells. In BCR-ABL1^+^ cells, SHP2 suppresses *Mxd3* and *Mxd4* through ERK and SFKs, likely leading to induction of select MYC targets, whereas in WT cells, SHP2 does not regulate the transcription of these genes. Our results suggest that SHP2 might be an alternative/additional target in BCR-ABL1-induced malignancies, a prospect made tangible with the recent development of bioavailable and highly selective SHP2 inhibitor.^59^

## Authorship Contributions

S.G. and B.G.N. designed the experiments. S.G., G.C., and W.Y. performed the experiments. S.G., A.Z., Z.L., and C.V. analyzed RNA-seq data. R.A.V. provided essential advice. S.G. and B.G.N. wrote and all authors edited the manuscript.

## Acknowledgements

We thank the Princess Margaret Genomics Centre for RNA-seq, and the Sick Kids-UHN Flow Cytometry Facility for FACS. This work was supported by R35CA49152 to B.G.N., and NIH R01 CA090576 to R.A.V.. B.G.N. was a Canada Research Chair, Tier 1, and work in his laboratory was partially supported by the Princess Margaret Cancer Foundation and a grant from the Ontario Ministry of Health and Long Term Care.

